# Unsupervised Clustering Reveals Spatially Varying Single Neuronal Firing Patterns in the Subthalamic Nucleus of Patients with Parkinson’s Disease

**DOI:** 10.1101/863464

**Authors:** Heet Kaku, Musa Ozturk, Ashwin Viswanathan, Joohi Shahed, Sameer Anil Sheth, Suneel Kumar, Nuri F. Ince

**Author notes:** Corresponding author: (Nuri F. Ince).

## Abstract

**Introduction:** Subthalamic nucleus (STN) is an effective target for deep brain stimulation (DBS) to reduce the motor symptoms of Parkinson’s disease (PD). It is important to identify firing patterns within the structure for a better understanding of the electro-pathophysiology of the disease. Using recently established metrics, our study aims to autonomously identify the discharge patterns of individual cells and examine their spatial distribution within the STN.

**Methods:** We recorded single unit activity (SUA) from 12 awake PD patients undergoing a standard clinical DBS surgery. Three extracted features from raw SUA (local variation, bursting index and prominence of peak) were used with k-means clustering to achieve the aforementioned unsupervised grouping of firing patterns.

**Results:** 279 neurons were isolated and four distinct firing patterns were identified across patients: tonic (11%), irregular (55%), periodic (9%) and non-periodic bursts (25%). The mean firing rates for irregular discharges were significantly lower (p<0.05) than the rest. Tonic firings were significantly ventral (p<0.05) while periodic (p<0.05) and non-periodic (p<0.01) bursts were dorsal. The percentage of periodically bursting neurons in dorsal region and entire STN were significantly correlated with off state UPDRS tremor scores (r=0.51, p=0.04) and improvement in bradykinesia and rigidity (r=0.57, p=0.02) respectively.

**Conclusion:** Strengthening the application of unsupervised clustering for firing patterns of individual cells, this study shows a unique spatial affinity of tonic activity towards the ventral and bursting activity towards the dorsal region of STN in PD patients. This spatial preference, together with the correlation of clinical scores, can provide a clue towards understanding Parkinsonian symptom generation.

## Introduction

The subthalamic nucleus (STN) is an established target in deep brain stimulation (DBS) to control the motor symptoms of Parkinson’s disease (PD). It has been suggested that firing pattern changes in single unit activity (SUA) within the basal ganglia could be a key in understanding the pathophysiology of PD [1] and can assist in electrophysiological mapping of the structure [2]. Studies involving 1-methyl-4-phenyl-1,2,3,6-tetrahydropyridine (MPTP) treated non-human primates report an increase in oscillatory neuronal activity and instantaneous firing rates [3] along with an increase in bursting activity [4]. Based on these observations, Wichmann and DeLong suggested that both, the change in firing rate and firing patterns are relevant to the pathophysiology of PD and investigating them have the potential to unravel the underlying neural network inducing PD manifestations [5]. In this regard, various groups have reported different types of neuronal firing patterns in the STN of Parkinsonian patients [2, 3, 6, 7]. In particular, oscillatory firings were separated from non-oscillating ones in [7], whereas others have focused on identification of tonic (regular), irregular and oscillatory firing patterns [6]. The majority of these studies characterized firing profiles in a binary fashion. The metrics used to separate these profiles are modestly unclear and, in several instances, chosen arbitrarily based on visual inspection such as inter-spike interval (ISI) histograms or binary spike trains. Notwithstanding, in total, three main firing patterns have been commonly described: tonic, irregular and oscillatory [2, 3, 6, 7]. Irregular firings are most common in proportion, followed by tonic and then oscillatory [2, 6]. Furthermore, studies also investigated the spatial distribution of oscillatory firing patters [7] as the dorsolateral STN is known to harbor the motor circuitry of STN, whereas the ventromedial STN is associated with limbic circuitry [6].

Grouping of firing patterns into different categories manually is difficult as it is both time consuming and, in some cases, hard to achieve using the limited information provided by an auditory/visual inspection. By using recently established metrics [8, 9], our study aims to automatically characterize the proportion and spatial distribution of four firing patterns: tonic (T), irregular (I), periodic bursts (PB) and non-periodic bursts (NPB) within the STN of PD patients revealing distinct firing profiles in the territories of STN.

## Materials & Methods

### 2.1 Patients and Recordings

The study was approved by the Institutional Review Boards at the University of Houston and Baylor College of Medicine. Intraoperative SUA was recorded from 12 awake PD patients (8 Male, 4 Female, Age: 54±11.14 years) under local anesthesia as part of the standard clinical procedure. Eight of the 12 patients underwent bilateral DBS implantation, and 4 had unilateral implants constituting 20 STN recordings in total. Patients were asked to stop medication at least 12 hours prior to the surgery. Three microelectrodes (NeuroProbe; AlphaOmega, Israel) were initially placed at least 15mm above the stereotactic target and advanced towards the target depth with 0.5mm steps. SUA was recorded from three micro-electrode trajectories using a bio-amplifier (Grapevine Neural Interface and Processor; Ripple, UT) at 30 KHz. During the awake surgery, real time SUA traces were displayed on a data-scope to help with the localization of the target structure. Entry and exit of the STN was determined by a clinical neurophysiologist by listening to and visually observing the firing patterns of neurons. The dorsal borders of the STN in each patient were defined by inspection of SUA showing an increase in background activity and discharge rates. The dorsal border was taken as reference and marked as 0mm with negative depth values ventral to it. Spatially, 0mm to −2.5mm was considered as the dorsal half of STN as previous studies show dominance of oscillatory activity in this area [7]. Other firings captured above or below the electrophysiologically defined borders of the STN were not included in our report. The de-identified data were transferred to a computer for offline processing.

### 2.3 Data Analysis

The neural data was analyzed offline in Matlab 2017b (Mathworks, MA) using custom scripts and a publicly available spike sorting toolbox [10]. Since the neurons are observed to display a non-stationary firing behavior over time [2], we analyzed and clustered the raw segments from all 3 tracks in 5s intervals with a 2.5s overlap. This procedure helped us in capturing the neurons’ non-stationary behavior and label each segment separately based on its firing characteristic. Consequently, rather than assigning each neuron to a single category such as tonic or irregular, we clustered the segments and assigned them into categories.

Following the high pass filtering of raw SUA data over 300Hz, spike detection and sorting were performed automatically by using WaveClus [10], an open-source Matlab toolbox. Segments with a firing frequency of less than 10 Hz and duration less than 3s were discarded. Three features: (i) local variation [8]; (*LV*), (ii) bursting index [9]; (*BI*) and (iii) prominence of peak (*P*) were used together with k-means [11] to group the spike trains. To analyze the spatial distribution of discovered clusters statistically, Wilcoxon sign rank test was performed within each firing pattern to check if the pattern was significantly dorsal or ventral and to assess the difference in firing rates between these patterns as well as the intensity of bursting (*BI*) and regularity (*LV*) in dorsal and ventral regions. Correlation with Unified Parkinson’s Disease Rating Scale (UPDRS) was done using Spearman rank correlation due to non-normality of the data. Three patients were excluded from the correlation analysis since they were external referrals and access to their detailed medical records and itemized UPDRS scores were limited (see supplementary Table S1).

#### 2.3.1 Local Variation (LV)

*LV* was used as a measure of regularity in firings and defined as:

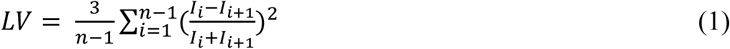

where I_i_ and I_i+1_ are the i^th^ and i+1^st^ ISI. *LV* helped eliminate the issues caused by variable firing rates and non-stationarity [8]. LV approaches 0 for a tonic firing neuron and 1 for a highly irregularly firing neuron.

#### 2.3.2 Bursting Index (BI)

Bursts were detected using a modified version of Max Interval Method [9]. For burst initiation, ISI below 10ms and a minimum of three consecutive spikes were required along with an intra-burst pause period of at least 20ms or a sum of 30ms between the next two ISIs. The bursting index is accepted as the ratio of the number spikes bursting to the number of spikes with ISI greater than 10ms.

#### 2.3.3 Prominence of Peak (P)

Upon initial visual investigation of the raw SUA data we observed that bursting can be both periodic and non-periodic. In order to distinguish between periodic and non-periodic bursts, a new feature called prominence of peak was computed from the spike train power spectrum. The digital spike train was convolved with a 10ms Gaussian window and its power spectral density (PSD) then calculated using Welch periodogram [12]. The peak value of the PSD was detected between 3 and 40Hz and its prominence was calculated using:

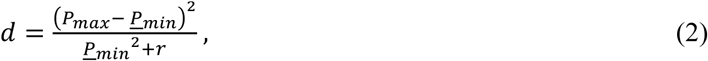

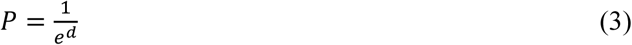

where *P* is the prominence, *P*_*max*_ is the peak value of PSD and *<u>P</u>*_*min*_ is the mean of the minimum values around this peak within an 8Hz bandwidth. A regularization parameter (r=10^−3^) was used to eliminate small peaks that might appear prominent due to a small background. Equation (3) was used to limit the prominence between 0 and 1 where values approaching 0 indicate strong oscillatory activity.

#### 2.3.4 Clustering

Clustering of firings into four categories was done using the k-means method [11] in a 3D feature space. The features were normalized before being subjected to the clustering algorithm. We varied the number of clusters from 1 to 8 and computed the sum of within-cluster distances for each index. When a relatively prominent elbow was observed in the graph, we used that index to decide on the total number of clusters in the data.

## Results

We analyzed 6705s of data and successfully isolated 279 neurons (23±7 neurons per patient). After spike detection and sorting, neurons were grouped into firing patterns and mapped onto the STN to investigate their spatial distribution. Figure 1A shows the border and depth definitions as well as exemplary raw SUA recordings from various depths. Figure 1B displays a representative trace for each irregular, tonic, periodic and non-periodic bursting firing pattern we studied. The clustering performed using the three features provided a prominent elbow around four for the total sum of intra-cluster spacing suggesting four distinct firing patterns (Figure 1C). To provide an example of their content, we provide 3 raw traces from each cluster as well. We observed that the largest cluster corresponding to 55% of the neuronal spiking displayed irregular patterns, where the remaining 11% displayed a tonic pattern while other 34% were bursting with 25% non-periodically and 9% periodically (Figure 1D). The mean firing rates for irregular firings (33.48±17.7 Hz) were significantly lower (p<0.05) than the tonic firings (74.19±33 Hz), periodic (82.35±43.46 Hz) and non-periodic bursts (57.57±28.66 Hz). Visual observation of the spatial distribution of these firing profiles (Figure 1E) revealed that the irregular firings were uniformly distributed throughout the STN. Interestingly, burst type firings populated the dorsal STN whereas tonic ones were tended towards the ventral end. We observed that the amount of tonic activity rapidly increased 3mm below the dorsal border of STN and extended towards the ventral region. Statistical analyses concluded that the tonic firings were significantly ventral (p<0.05) while periodic (p<0.05) and non-periodic bursts (p<0.01) were significantly dorsal. However, it should be noted that bursting patterns can still be observed within the ventral territory of the STN. Similarly, our findings show that both bursting (BI: 0.86±0.17 vs 0.79±0.2) and regularity (LV: 0.31±0.06 vs 0.21±0.08) were significantly different in the two regions of STN (p<0.01) with bursting being more intense in the dorsal region and tonic discharges being more regular in the ventral region. Finally, as done in [7] we correlated the off drug UPDRS part III scores and their improvement with medication with firing characteristics and found that the percentage of dorsally located periodic bursting neurons was significantly correlated with tremor (r=0.51, p=0.044). The percentage of periodic bursting neurons in the entire STN was significantly correlated with the improvement in bradykinesia and rigidity (r=0.57, p=0.02). Other UPDRS sub-scores did not show any significant correlation (p>0.05).

**Figure 1.**
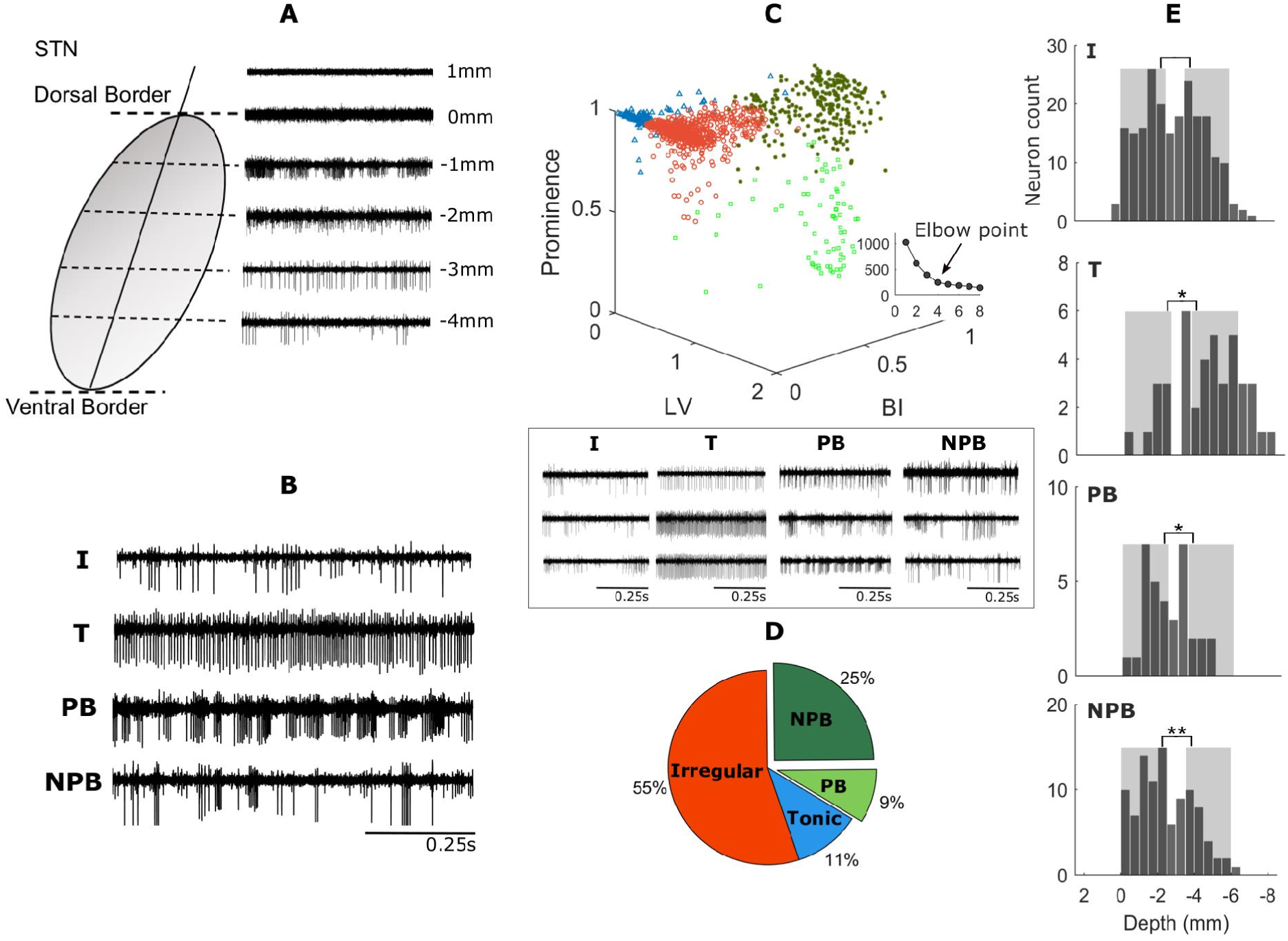
(A) Representation of STN and raw SUA recordings from various depths (0mm is the reference dorsal border). (B) Exemplary raw SUA data for four firing patterns (I=Irregular, T=Tonic, PB=Periodic burst, NPB=Non-periodic burst). (C) Clustering result over 12 subjects (1371 segments) (LV=Local Variation, BI=Bursting Index). Inset - Inter cluster distance plots for 1 to 8 clusters (x-axis: number of clusters, y-axis: sum of inter-cluster distance) and three examples of raw SUA traces from each cluster. (D) Proportion of firing patterns distributed across 12 subjects. (E) Spatial distribution of firing patterns across STN. The shaded areas indicate dorsal (0 to −2.5mm) and ventral (−3.5 to 6mm) regions of STN. (* indicates significant difference between shaded regions) (* = p < 0.05, ** = p < 0.01)

## Discussion

In this study, using an unsupervised approach utilizing recently established metrics, we identified and investigated the spatial distribution of neurons within the STN of PD patients based on their firing patterns. In addition to commonly reported irregular, tonic and bursting patterns [2, 6], we further divided the bursting category into oscillatory and non-oscillatory bursts. While most studies compare the spatial distribution of oscillatory units against non-oscillatory units [7], we expanded the investigation by looking at the spatial distribution of each of the four individual patterns separately. An interesting and novel finding was the occurrence of tonic firings towards the ventral region of STN. While the presence of oscillatory neurons in the dorsal region has already been established [7], our study shows the occurrence of burst type of findings (be it periodic or non-periodic) in the dorsal region as well. Our observations of irregular firings dominating the set (55%), followed by non-periodic bursts (25%), tonic (11%) and periodic burst firings (9%) were comparable to previous findings [2, 3]. The high fraction of bursting in our findings is also in accordance with the previously studied MPTP treated animal models [3, 4] where the main difference between healthy and Parkinsonian state was the emergence of bursting activity and increased instantaneous firing rates. Similar to the study conducted by Weinberger et al. [7], we found a significant correlation between the percentage of periodically bursting dorsal neurons with the off state tremor score and overall percentage of periodic neurons with improvement in bradykinesia. Given the somatotopic organization of STN [6], our observation of the predominance of bursting activity in dorsal STN and tonic activity in ventral region could be indicative of variable disease manifestations such as motor and cognitive/limbic features of PD. The study of pattern changes as well as their distribution within the structure thus could provide a better understanding of the more complex mechanisms underlying the disease.

We presented an automated approach that can isolate clusters of neurons based on their firing profiles in the territories of STN. In practice 83% of centers around the world are using MER for electrophysiological mapping to refine the final position of the chronic electrode [13]. Intraoperative decision making involves the conversion of single-unit neuronal activity recorded at the microelectrode tip, into audio and visual signals. This process of mapping is time consuming experience-based and depends critically on the ability of neurosurgeons or electrophysiologists to recognize a variety of cues in the recorded neural activity. Multiple microelectrodes isolating neurons with distinct firing patterns makes the interpretation of auditory cues even more challenging. We propose that the presented automated approach can be used to investigate the firing profiles of a large number of neurons recorded simultaneously during DBS surgery. We postulate that it may have utility in identifying the most favorable track for macroelectrode implantation, and contact localization (i.e. depth), while also providing additional insight regarding the relationship of disease symptoms to different territories of the STN. Since the bursting firings were biased towards the dorsal region, the track with larger number of bursting neurons can be utilized for chronic DBS electrode placement. Nevertheless, further research will be required to better define how this technique can be applied to improve targeting of the dorsolateral STN where the majority of patients will experience clinical improvement.

It must be noted that using existing clinical electrodes one can record only 1-2 neuron at best per track at each depth thus limiting the number of neurons isolated within the territories of STN of each patient. We anticipate that, an increased patient population as well as future clinical electrodes designs with multiple SUA recording contacts can improve the sampling capacity and will likely provide advantages in establishing a bridge between single neuronal activity and clinical observations.

## Supplementary Information

### Surgical Protocol

Surgeries were performed in awake patients using local anesthesia. Patients were asked to discontinue Parkinson’s medications 24 h before DBS surgery. IV anesthetic agents were used at a low level during the placement of the head frame until the burr holes are placed. The anesthetic was stopped just prior to placement of the burr holes and recordings were commenced when the patient was alert and cooperative. This is usually 20-30 minutes after cessation of the anesthetic. The radiological coordinates and trajectories to the STN were identified by fusing preoperative stereotactic MRIs to a preoperative or an intraoperative stereotactic computed tomography scan on a neuro-navigational platform (StealthStation; Medtronic). Using a BenGun configuration, starting from 15 mm above the radiographic target, at least 15-20s of recordings were obtained at each depth until one of the microelectrode tips had first passed through the dorsal and then the ventral border of STN. Electrodes were moved using a 0.5-1mm step size to allow more precise identification of the STN borders. The location of the dorsal entry and ventral exit from the STN was determined by an experienced clinical neurophysiologist through listening to and visually observing the microelectrode recording of single-unit activities (MER-SUAs). As others have described^1,2^, the dorsal border of STN is detected when a prominent increase in background activity or uniform firing characteristic of border cells are observed. The location of the dorsal border of STN is used as reference depth (0 mm), and the characteristics of SUA patterns in dorsal and ventral regions of STN are reported in terms of relative distance to the border. If a track with this typical span was identified, and clinical benefit during stimulation testing was observed, the targeting was characterized as accurate. Generally, one of the tracks had a larger spatial span of neural activity. Additionally, there were other firings which were captured above or below the ventral borders of the STN likely due to electrode passing through thalamus and SNr. However, we only used SUA data between the dorsal and ventral borders.

**Table S1:** Patient Demographics. The total score was the sum of all 33 items in the MDS-UPDRS part III section. Tremor subscores (max. 16) were calculated for the right hemibody as the sum of ‘Postural Tremor-Right Hand’, ‘Kinetic Tremor -Right Hand’, ‘Rest Tremor -RUE’, ‘Rest Tremor -RLE’. PIGD subscores (max. 20) were calculated as the sum of ‘Arising From Chair’, ‘Gait’, ‘Freezing Of Gait’, ‘Postural Stability’, ‘Posture’. Bradykinesia+Rigidity subscores (max. 28) were calculated for the right hemibody as the sum of ‘Rigidity -RUE’, ‘Rigidity -RLE’, ‘Fingertap -Right Hand’, ‘Hand Movements -Right Hand’, ‘Pronation - Right Hand’, ‘Toe Tap –Right Foot’, ‘Leg agility -Right Leg’^3^. Three patients with missing UPDRS scores were externally referred and access to their detailed medical records was limited.

| Pt # | Sex | Age | Disease Duration | LEDD | Hemi-sphere | OFF Tremor Score | OFF PIGD Score | OFF bradykinesia +Rigidity Score | Total OFF PartIII Score | ON Tremor Score | ON PIGD Score | ON Akinesia Score | Total ON PartIII Score |
| --- | --- | --- | --- | --- | --- | --- | --- | --- | --- | --- | --- | --- | --- |
| 1 | M | 53 | 9 | 599 | Right | 5 | 2 | 12 | 33 | 4 | 1 | 9 | 24 |
| 2 | F | 50 | 8 | 700 | Left | 2 | 2 | 6 | 26 | 0 | 2 | 2 | 12 |
| 3 | F | 62 | 5 | 683 | Left | 2 | 2 | 0 | 18 | 0 | 0 | 0 | 7 |
| 4 | F | 49 | 6 | 1464 | Left | 4 | 2 | 17 | 56 | 1 | 0 | 10 | 27 |
| 4 | F | 49 | 6 | 1464 | Right | 5 | 2 | 19 | 56 | 2 | 0 | 10 | 27 |
| 5 | F | 63 | 12 | 940 | Left | 1 | 5 | 16 | 46 | 1 | 3 | 12 | 33 |
| 5 | F | 63 | 12 | 940 | Right | 2 | 5 | 13 | 46 | 1 | 3 | 11 | 33 |
| 6 | M | 41 | 7 | 1375 | Left | 2 | 4 | 16 | 57 | 0 | 2 | 1 | 13 |
| 6 | M | 41 | 7 | 1375 | Right | 5 | 4 | 18 | 57 | 0 | 2 | 6 | 13 |
| 7 | M | 40 | 8 | 1103 | Left | 3 | 4 | 11 | 45 | 0 | 1 | 6 | 23 |
| 7 | M | 40 | 8 | 1103 | Right | 2 | 4 | 16 | 45 | 1 | 1 | 13 | 23 |
| 8 | M | 58 | 8 | 873 | Left | 3 | 2 | 17 | 48 | 0 | 0 | 6 | 16 |
| 8 | M | 58 | 8 | 873 | Right | 2 | 2 | 15 | 48 | 0 | 0 | 5 | 16 |
| 9 | M | 64 | 9 | 1300 | Left | 3 | 3 | 10 | 32 | 3 | 1 | 10 | 23 |
| 9 | M | 64 | 9 | 1300 | Right | 2 | 3 | 7 | 32 | 2 | 1 | 2 | 23 |
| 10 | M | 63 | 7 | 900 | Left | 9 | 4 | 10 | 47 | 8 | 3 | 6 | 32 |
| 10 | M | 63 | 7 | 900 | Right |  |  |  |  |  |  |  |  |
| 11 | M | 35 | 6 |  | Right |  |  |  |  |  |  |  |  |
| 12 | M | 70 |  |  | Left |  |  |  |  |  |  |  |  |
| 12 | M | 70 |  |  | Right |  |  |  |  |  |  |  |  |

LEDD: Levodopa equivalent daily dose<sup>4,5</sup>

**Figure S1.**
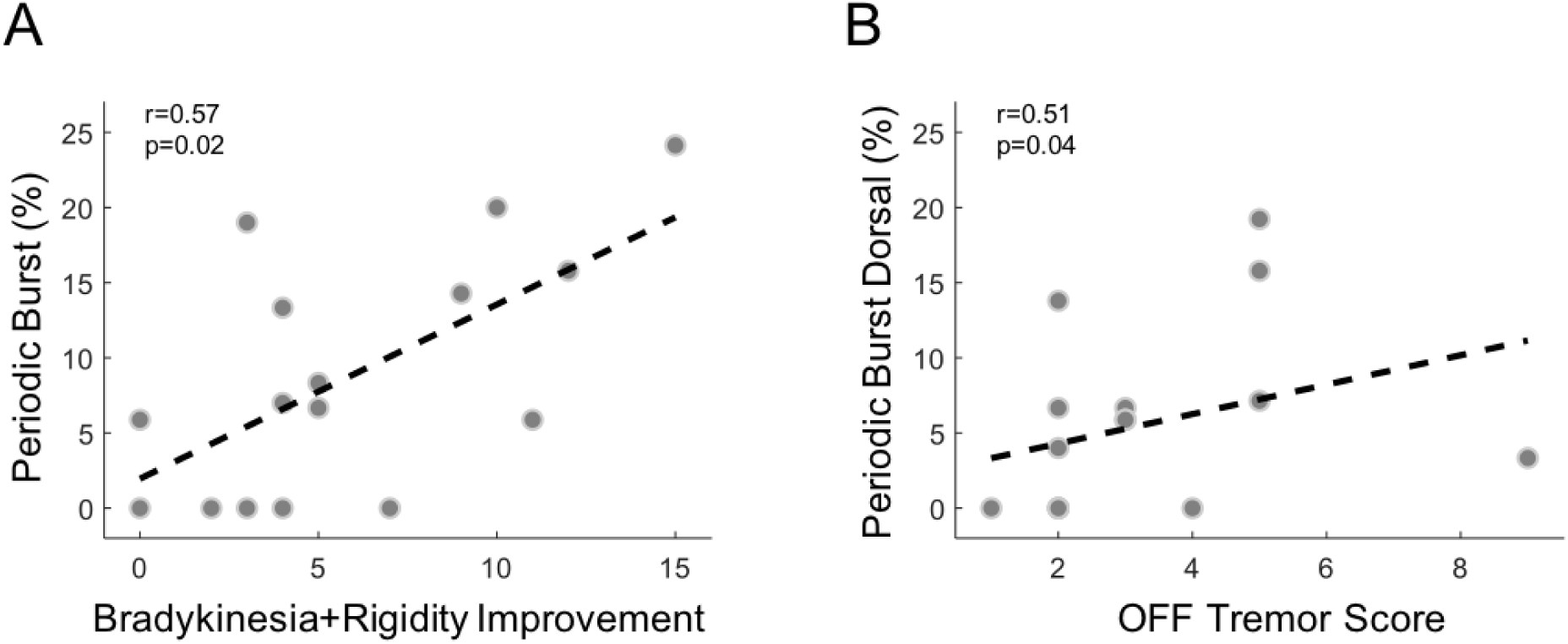
Scatter plots for correlation between (A) percentage of periodic bursting neurons and bradykinesia & rigidity improvement and (B) percentage of periodic bursting neurons found dorsally within STN with OFF state tremor scores. (n=16 for both plots). The analyses were performed between the hemibody UPDRS and SUA from the contralateral STN and were controlled for age, sex and disease duration by fitting a linear model with the interaction terms to the data. The UPDRS measure was provided as response variable and neural measure was provided as predictor alongside with age, sex (as a categorical variable) and disease duration and their interaction terms. Our analyses indicated that the UPDRS measures did not differ significantly according to age, sex or disease duration.

